# Thymidylate synthase drives the phenotypes of epithelial-to-mesenchymal transition in non-small cell lung cancer

**DOI:** 10.1101/2020.08.13.249581

**Authors:** Mohammad Aarif Siddiqui, Paradesi Naidu Gollavilli, Vignesh Ramesh, Beatrice Parma, Annemarie Schwab, Maria Eleni Vazakidou, Ramakrishnan Natesan, Ozge Saatci, Ida Rapa, Paolo Bironzo, Harald Schuhwerk, Irfan Ahmed Asangani, Ozgur Sahin, Marco Volante, Paolo Ceppi

## Abstract

**Background:** Epithelial-to-mesenchymal transition (EMT) enhances motility, stemness and chemoresistance of carcinomas and is an important determinant of metastasis. Little is known about how various pathways coordinate to elicit EMT’s different functional aspects. Even lesser has been studied in this context in non-small cell lung cancer (NSCLC), where EMT is a key event during early tumorigenesis. Thymidylate synthase (TS), a proliferation enzyme, has been previously correlated with EMT transcription factor ZEB1 in NSCLC and is associated with resistance against anti-folate chemotherapy. In this study we establish a functional correlation between TS, EMT, chemotherapy and metastasis and identify a network that might propel the TS mediated EMT phenotype.

**Methods:** Published datasets were analysed to evaluate significance of TS in NSCLC fitness and prognosis. mCherry based promoter reporter assay was used to sort Calu-1 and A549 NSCLC cells in TS^HIGH^ and TS^LOW^. Metastatic potential of TS knock-down was assayed in syngeneic C57BL/6 mice.

**Results:** Low TS levels were prognostic and predicted chemotherapy response. NSCLC cell lines with higher TS promoter activity were more mesenchymal-like. RNA-seq from these cells, and shRNA mediated TS knocked down cells, identified EMT as one of the most differentially regulated pathways. EMT transcription factors HOXC6 and HMGA2 were identified as upstream regulator of TS whereas, AXL, SPARC and FOSL1 were identified as downstream effectors. TS knock-down reduced the metastatic colonisation *in vivo*.

**Conclusion:** These results establish the role of TS as a theranostic NSCLC marker integrating survival, chemo-resistance and EMT, and identifies a regulatory network that could be exploited to target EMT-driven NSCLC.

## Introduction

Epithelial-to-mesenchymal transition (EMT) is an embryonic process hijacked by epithelial-like carcinoma cells to gain mesenchymal-like phenotype. Oncogenic EMT is a gamut of functional changes, such as enhanced motility, invasiveness, stemness, aggressiveness and chemoresistance, and is a key determinant of metastasis. EMT is complex cascade of molecular events engendered by master EMT transcription factors (EMT-TFs, ZEB1/2, SNAI1/2 and TWIST) in response to extracellular cues including cytokines and hypoxia(1, 2). EMT-TFs activate multiple molecular pathways that ultimately leads to alteration in cytoskeleton and cell-adhesion proteins(3). EMT is a key early event in NSCLC biology and steers epithelial-like cells towards stemness, chemoresistance and metastatic dissemination(4, 5). It is engineered through coordination of divergent molecular pathways(6), and presumably orchestrated by different EMT-TFs at different progression time points(7–9). How these pathways connect to each other and affect different modalities of EMT is still largely unexplored.

Thymidylate synthase (TS) is a *de novo* pyrimidine biosynthesis enzyme that catalyzes conversion of deoxyuridine monophosphate to thymidine monophosphate, essential for DNA synthesis and cell proliferation. It is targeted by chemotherapeutic drugs, like pemetrexed, in NSCLC, and has been widely studied as a chemoresistance marker(10). Our lab recently showed a correlation between expression of TS and EMT markers in NCI-60 panel of cancer cell lines originating from different tissues(11), and established its role in maintaining the de-differentiated mesenchymal-like state of triple negative breast cancer(12). In this study we present evidence that TS is not a mere proliferation marker in NSCLC, but also has a direct role in driving EMT phenotypes, with several biological and clinical implications.

## Methods

### Cell lines

A549 (NCI) and Calu-1 (ATCC) were cultured in RMPI-1640, supplemented with 10% FBS, 1%Pen/Strep and 1%L-Glutamine (all from Sigma). LL/2, Ladi 3.1 and Ladi 2.1 cells were cultured in DMEM (Sigma) supplemented with 10% FBS, 1%Pen/Strep and 1%L-Glutamine. Human cells were STR-profiled, used between passages 3 and 15, examined for mycoplasma and maintained in Plasmocin (Invivogen) to prevent contamination.

### Lentiviral transduction

Plasmids for TS knock down (TRCN0000456666) is from Sigma. Empty backbone pLKO.1 (referred to as pLKO) was used as control. Plasmids from *TYMS*-promoter reporter (HPRM33357-LvPM02), GAPDH promoter reporter (HPRM39787-LvPM02), TS expression vector (Ex-T0406-LV105b) and control vector (Ex-Neg-LV105b) are from GeneCopoeia. For production of lentiviral particles, 293T cells were transfected with 8μg knock-down/expression vectors and 2μg of pMDL, pVsVg and pRevRes in complex with 24μg PEI (Polysciences). After 48 hours, supernatant was collected, centrifuged and filtered. For transduction, 10^5^ cells were seeded in a 6-well plate and infected in presence of 8μg/ml polybrene (Sigma). Selection was done with 3μg/ml puromycin (Sigma) and cells were maintained in 1 μg/ml puromycin.

### RNA sequencing

Total RNA was extracted using miRNeasy kit (Qiagen) following the manufacturer’s instructions. RNA-Seq libraries were constructed using the TruSeq sample Prep Kit V2 (Illumina). Briefly, 1μg of purified RNA was poly-A selected and fragmented with fragmentation enzyme. After first and second strand synthesis from a template of poly-A selected/fragmented RNA, other procedures from end-repair to PCR amplification were done according to library construction steps. Libraries were purified and validated for appropriate size on a 2100 Bioanalyzer High Sensitivity DNA chip (Agilent Technologies.). The DNA library was quantified using Qubit and normalized to 4nM before pooling. Libraries were pooled in an equimolar fashion and diluted to 10pM. Library pools were clustered and run on Nextseq500 platform with paired-end reads of 75 bases, according to the manufacturer’s recommended protocol (Illumina). Raw reads passing the Illumina RTA quality filter were pre-processed using FASTQC for sequencing base quality control. Sequence reads were mapped to UCSC human genome build using TopHat and differential gene expression determined using Cufflinks 2.1.1 and Cuffdiff2.1.1 as implemented in BaseSpace. The sequencing data has been submitted GEO dataset and could be accessed with GSE148589 accession number.

### Quantitative real-time PCR

Total RNA was extracted using miRNeasy kit (Qiagen) and 50ng was converted to cDNA using Tetro cDNA synthesis kit (Bioline) with random hexamers. GAPDH was used as the internal control. TaqMan probes (Thermo-Fisher) were used for quantification in Applied Biosystems 7300. Fold change was calculated using the ΔΔCt method.

### Gene Set Enrichment Analysis

Gene set enrichment analysis (GSEA), for computing overlap, on the differentially expressed genes upon TS knockdown was performed with the geneset collections in the Molecular Signatures Database v6.1 software. For EMT geneset enrichment analysis in the patient data, normalized gene expression values were downloaded from GEO database (GSE101929) and cbioportal platform for TCGA profile (LUAD, PanCaner). For calculation of TS Knockdown (KD) score, first, z scores of the down- and up-regulated genes upon TS knockdown were calculated. Then, the sum of z scores of downregulated genes was subtracted from the sum of z scores of upregulated genes and KD scores were obtained for each patient. Patients were grouped for the analysis based on either the median value of TYMS gene expression or KD score.

### Survival Analysis

Normalized gene expression profiles of lung cancer samples were downloaded from GEO (GSE50081, GSE72094, GSE30219) and mRNA expression values as Z-scores were obtained for TCGA profiles (LUAD and LUSC) from cbioportal platform. Thirty-five samples from completely resected NSCLC patients were collected from the files of San Luigi Hospital, Orbassano, Turin, Italy. None of the patients received either neo-adjuvant chemotherapy or radiation therapy and all received adjuvant cisplatin and pemetrexed. All cases were reviewed and classified using anonymized samples. Clinical samples were stratified as TYMS-low and TYMS-high based on the median value of gene expression as cut-off. Kaplan-Meier estimate was used to generate survival curves and significance between the two groups were analyzed using log-rank test in R software. Survival graphs from the KM Plotter database was generated based on TYMS expression by using the auto select best cut-off option. TS KD score for survival curve was calculated as described in the previous section.

### Western blot analysis

Cells were lysed in RIPA buffer and quantified using Pierce BCA kit (Thermo-Fisher). Proteins lysates (10 – 20μg) were resolved on 10% SDS–PAGE gels and transferred to PVDF membrane (Thermo-Fisher). Membranes were blocked in 5% Milk (BioRad) in 1XTBS-T and incubated overnight in primary antibodies diluted in 5% milk at 4°C. anti-TS (EPR4545) and -SPARC (SP205) antibodies were purchased from Abcam; anti-E-Cadherin (4A2), -Vimentin (D21H3), -AXL (C89E7), -FOSL1 (D80B4) and -β-Actin (8H10D10) were purchased from Cell Signaling. After incubation with secondary antibodies (Southern Biotech), detection was performed using the ECL (Thermo-Fisher) and developed on X-Ray film (Thermo-Fisher) using a chemiluminescence imager, AGFA CP100.

### Proliferation assay

For proliferation assay cells were seeded in 96-well plates in low density (5-20% initial confluency). Plates were loaded in Incucyte-Zoom (Essen Bioscience) and scanned every 2-4 hours. For each scan, phase contrast image was acquired from every well and was analyzed by Incucyte Zoom software.

### *in vitro* drug treatment

Pemetrexed was purchased from Sigma. For *in vitro* treatment cells were plated in a 96-well plates (4,000 cells/well) and incubated overnight. For cytotoxicity death assay, 2000X Cytotox Green Reagent (Essen Bioscience) was diluted in RPMI and working dilutions of pemetrexed was prepared in Cytotox Green supplemented media. After treatment, plate was loaded in Incycuyte Zoom and images were acquired in real-time for phase to quantify growth. Activity of Cytotox reagent was simultaneously acquired at the green channel to quantify death. Incycuyte Zoom software was used for the analysis and data export.

### Migration assay

For migration assay cells were plated in 96-well plates so that they reach 90% confluency overnight. Cells were wounded using WoundMaker (Essen Biosciences) as per the instruction from the manufacturer. Plates were loaded in Incucyte Zoom and were automatically scanned for programmed time interval. For each scan, wound width was recorded by the software and the proliferation inside the wound was normalized to the proliferation outside the wound, giving relative wound density for each time point.

### Tumorsphere culture

40,000 cells were seeded in triplicates in ultra-low attachment 6-well plates (Corning) in complete Mammocult medium (Stem Cell Technologies), prepared according to the manufacturer’s instruction. After formation, spheres were counted by spinning at 300g for 5 minutes and suspending in PBS (Lonza).

### siRNA transfection

Reverse transfection was done with Lipofectamine RNAiMAX Transfection Reagent (Thermo). 50nM siRNA were mixed with 1.5μl transfection reagent in 200μl Opti-MEM (Thermo) and incubated for 15 minutes. After incubation transfection complex was added to the bottom of 12-well plates and 10^5^ cells, suspended in 800μl, were added. Cells were incubated at 37°C, 5%CO_2_. Cells were lysed for western blot after 72 hours.

### Subcutaneous Injection

For subcutaneous injections, 1.0×10^6^ cells resuspended in 0.9% NaCl were mixed with Matrigel (Corning) in a ratio 1:1 (v:v). Cells were injected in right flanks of 10–15 weeks old female C57BL/6, with 8 mice per group. Caliper measurements were taken every 4th day and the tumor volume was calculated using the formula (Length*Width^2*π)/6.

### Tail-vein injection

For tail-vein metastasis assay, 5×10^5^ LL/2 pLKO and shTs cells resuspended in 100μl PBS and injected in tail vein of 8-10 weeks old female C57BL/6, with 10 mice per group. Lung metastases were monitored by bioluminescence imaging (BLI) 4 weeks after injection. Anesthetized mice were intraperitoneally injected with 150 mg/kg D-luciferin (Kayman Chemicals). Bioluminescence images were acquired with Lumina III in vivo Imaging System (IVIS, Perkin Elmer). For all the mice exposure time was maintained at 180 seconds. Raw IVIS images were analysed with Living Image software and the metastasis was represented as radiance.

### Statistical analysis

Statistical tests were performed with the GraphPad software v.7 comparing groups of different conditions with replicates. In all tests, the statistical significance was set at p=0.05 (in the figures * indicates p<0.05, ** p<0.01, *** p<0.001).

## Results

### TS is an essential NSCLC gene with prognostic/predictive power and correlates with EMT signatures

We evaluated different clinical aspects of TS in NSCLC and assayed its correlation with EMT. As a rate-limiting *de novo* pyrimidine biosynthesis enzyme, *TYMS* (gene coding TS) has been proposed as an essential gene, but, so far, no functional data have been shown in NSCLC. To evaluate dependency of NSCLC on TS, a dataset generated from a genome-wide CRISPR/Cas9 screen of 18,009 genes in 324 cancer cell lines was exploited(13). Based on a gene fitness score that defined how strongly a cancer is dependent on a gene for survival and growth, a priority score was generated to identify the most promising drug targets. Among all the pan-cancer fitness genes identified, TS ranked 31^st^ (top 1%, **Figure 1A**). In NSCLC subsets, it ranked 2^nd^ and 19^th^ in squamous cell carcinoma (SCC) and adenocarcinoma (ADC) respectively (**Figure 1B**), indicating that NSCLC strongly depend on TS for sustained growth. TS was also found to be 5^th^ among the pan-cancer priority targets identified (**Supp. Fig 1A)**. TS expression has been consistently found increased in NSCLC compared to adjacent normal tissues(14) and correlated with poor prognosis in different expression datasets (**Figures 1C-D**, **Supp. Figures 1B-C**). TS is targeted by the antifolate drug pemetrexed, and its overexpression has been proposed to determine chemoresistance(15). For *in vitro* validation, we established shRNA-mediated TS knockdown in two NSCLC cell lines and observed a significant increase in pemetrexed sensitivity (**Figures 1E-F**). To test if *in vitro* evidence also reflected in outcome of chemotherapy-treated patients, we retrospectively analyzed a small case-series of NSCLC patients treated with pemetrexed-based chemotherapy and found that higher TS gene expression significantly associated with worse prognosis (**Figure 1G**). These results emphasize the importance of TS as a prognostic and predictive marker, in line with previous literature(10). However, chemoresistance is also an important hallmark of EMT, and recent pivotal findings from our lab associated TS expression with EMT markers in cancers from different origins and suggested a potential direct role(11). To test this in NSCLC, we analyzed cells belonging to the CCLE dataset, categorized as epithelial or mesenchymal based on ratio of Vimentin (*VIM*) and E-Cadherin (*CDH1*) expression(16) (**Supp. Figure 1D**) and found that mesenchymal-like cells expressed higher TS compared to epithelial-like (**Figure 1H**). To further demonstrate its clinical significance, we investigated multiple datasets and found that patients with higher TS expression were significantly enriched for hallmark EMT genes (**Figures 1I-J**, **Supp. Figures 1E-F**).These results indicate that TS is not only an essential proliferation gene with a strong prognostic and predictive role, but also has a potential role in EMT in NSCLC.

**Figure 1.**
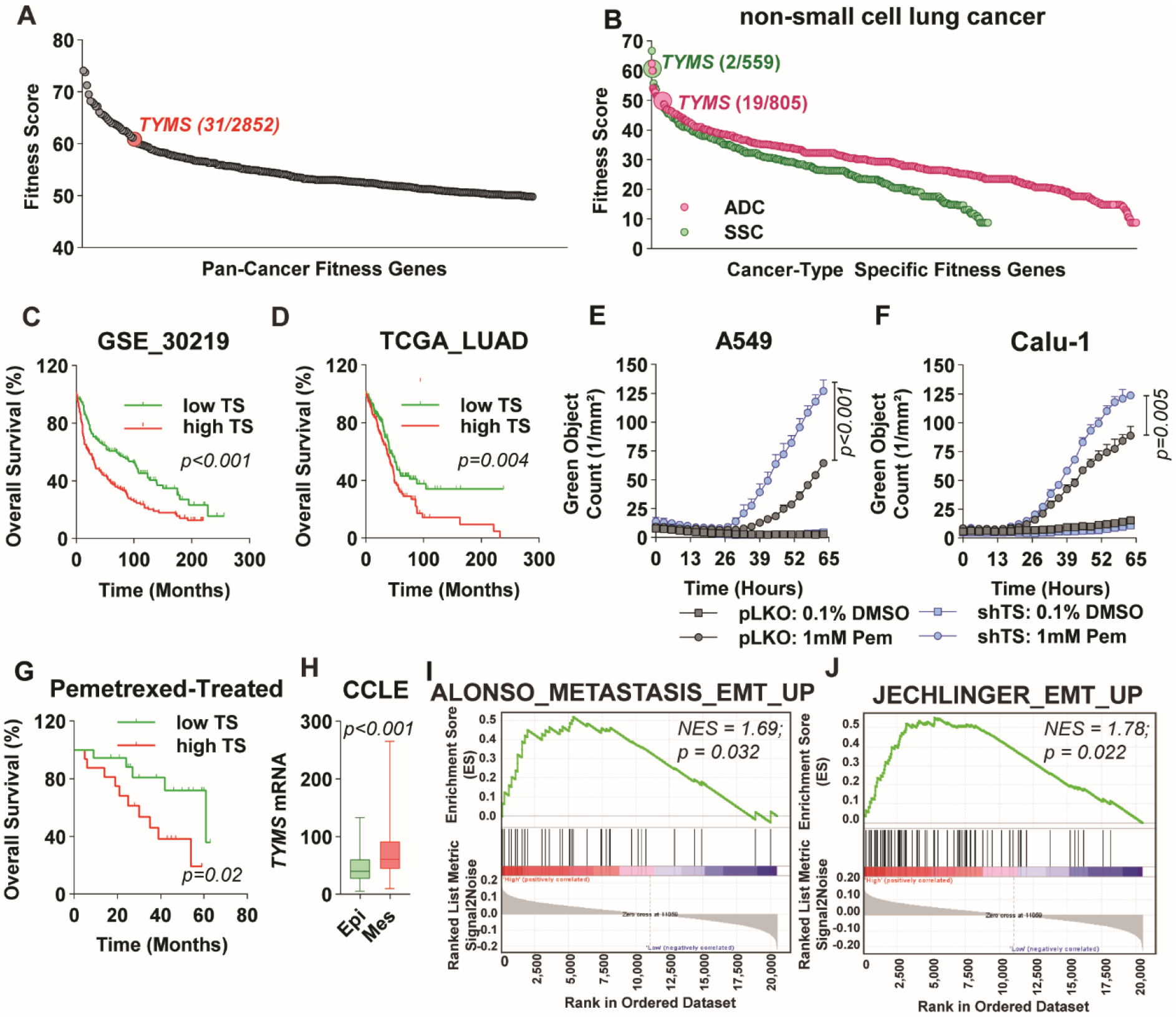
TS is an essential NSCLC gene with prognostic/predictive power and correlates with EMT signatures. (**A**) Graph showing TS ranking as a pan-cancer fitness gene for growth and survival. (**B**) Fitness score for TS in lung adenocarcinoma (ADC) and squamous cell carcinoma (SCC). (**C-D**) Predicted overall survival in non-small cell lung cancer (NSCLC) patients separated according to the expression level of TS. p-value is Logrank test. Sensitivity of NSCLC cells lines A549 (**E**) and Calu-1 (**F**) to anti-folate drug pemetrexed (Pem) after shRNA mediated knock-down of TS. Dead cells have been quantified as green object counts. p-value is represented as 2-way ANOVA, Sidak’s multiple test. (**G**) Kaplan-Meier survival analysis of NSCLC patients treated with pemetrexed-based chemotherapy categorised according to high and low TS mRNA expression. p-value is Logrank test (**H**) *TYMS* mRNA levels compared between NSCLC epithelial (Epi) and mesenchymal (Mes) cells from CCLE data defined by *VIM*/*CDH1* mRNA expression ratio. p-value is from a Student’s t-Test. (**I-J**) Gene-set enrichment analysis of indicated datasets with high and low TS correlating with EMT gene signature in TCGA lung adenocarcinoma data set. Error bars represent standard deviation.

### Endogenous TS level as an important determinant of EMT phenotype

We aimed at studying the role of intrinsic TS level on EMT *in vitro*, coupled with functional approaches. To do so, Calu-1, a NSCLC cell line, was stably transduced with a promoter reporter construct that expressed mCherry fluorescent protein transcribed from *TYMS* promoter (**Figure 2A**). After puromycin selection, cells were FACS-sorted for highest and lowest red fluorescence (indicated further as TS^HIGH^ and TS^LOW^) (**Figure 2B**). TS^LOW^ cells proliferated slower (**Figure 2C**) and showed a distinct epithelial phenotype, whereas TS^HIGH^ resembled a mesenchymal-like morphology (**Figure 2D).** To confirm differential EMT status at molecular level, expression of E-CAD and VIM, markers for epithelial-like and mesenchymal-like cells respectively, was quantified. At mRNA level, there was a striking difference between the expression of *CDH1* (gene coding E-CAD) and *VIM* (**Figure 2E)**. TS^LOW^ cells expressed more E-CAD and lesser VIM compared to TS^HIGH^ cells also at protein level (**Figure 2F)**. When assayed in proliferation-normalized wound migration assay, TS^LOW^ cells migrated slower than TS^HIGH^ cells (**Figures 2G-H)**. As a control for the promoter reporter assay, cells were sorted for GAPDH promoter activity and no difference in EMT markers and migration was observed in GAPDH^HIGH^ and GAPDH^LOW^ cells (**Figures 2F-H)**. TS^LOW^ cells also had reduced self-renewal capacity, quantified as the number of tumorspheres formed in a low-adherence culture (**Figure 2I**).

**Figure 2.**
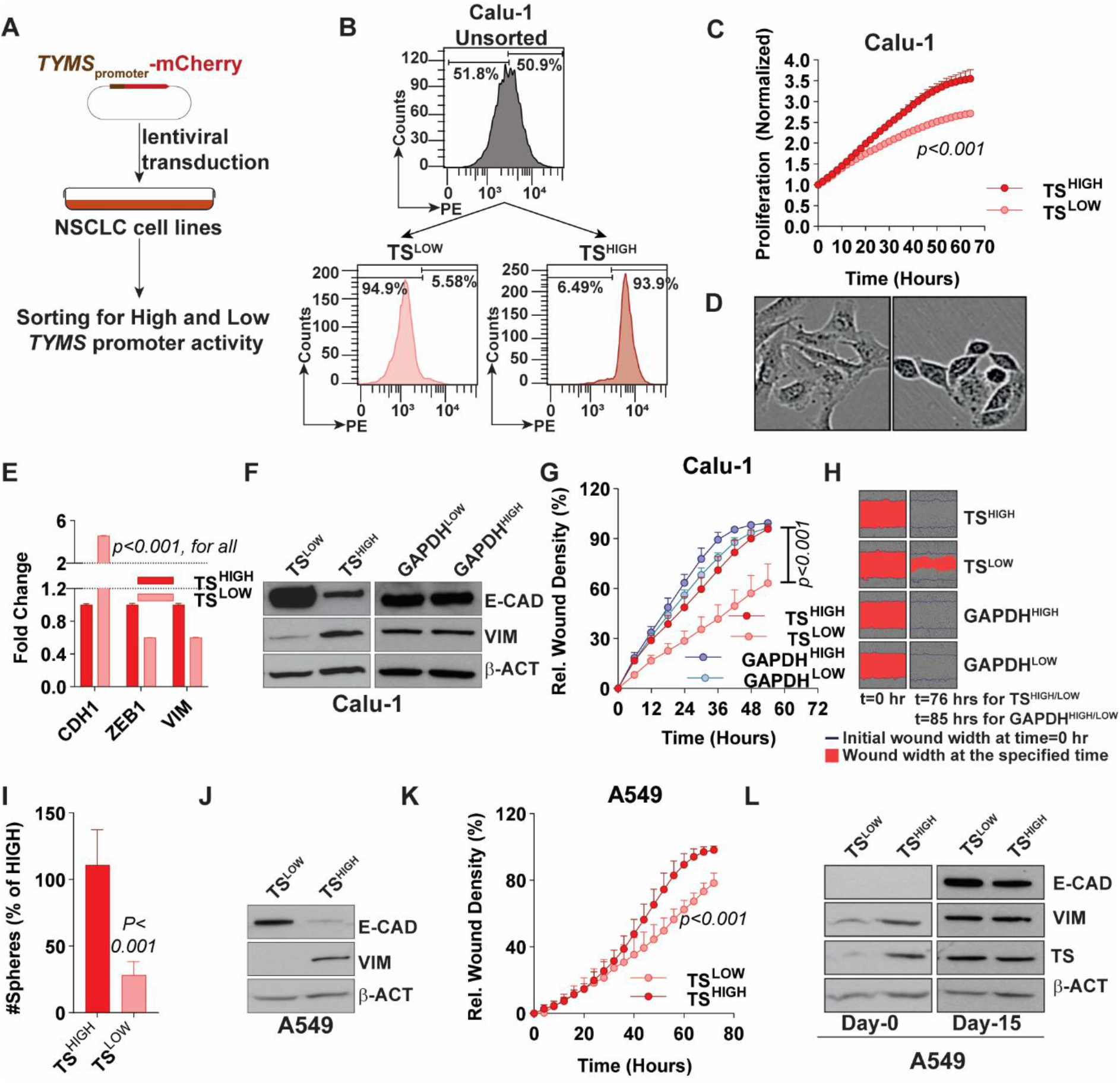
NSCLC cells can be sorted into distinct EMT phenotype based on *TYMS* promoter activity. (**A**) Calu-1 cells were stably transduced with a lentiviral vector expressing mCherry gene under control of *TYMS* promoter. Transduced cells were selected for puromycin positivity and 5% Calu-1 cells were sorted with highest and lowest expression of mCherry. (**B**) FACS plots depicting the separation of high and low population of Calu-1 cells from scheme presented in A. The numbers on the FACS plot represent the percentage of TS^HIGH^ and TS^LOW^ population in the unsorted cells. (**C**) Real-time proliferation curve showing difference in growth between Calu-1 TS^HIGH^ and TS^LOW^ cells. Cells have been normalized to confluency at time 0, to exclude the difference in surface areas. p-value is represented as 2-way ANOVA, Sidak’s multiple test. (**D**) Picture showing morphology of Calu-1 cells sorted for high *TYMS* promoter activity (TS^HIGH^) and low promoter activity (TS^LOW^). (**E**) mRNA quantification of *CDH1, ZEB1* and *VIM* in Calu-1 TS^HIGH^ and TS^LOW^ cells. p-value is represented by multiple t-Test. (**F**) Protein quantification of E-CAD (marker of epithelial phenotype) and VIM (marker of mesenchymal phenotype) in TS^HIGH^ and TS^LOW^ Calu-1 cells. Cells were sorted in GAPDH^HIGH^ and GAPDH^LOW^ based on the scheme depicted in A but with mCherry transcript driven by GAPDH promoter. GAPDH sorted cells were used as control. (**G**) Real-time proliferation-independent migration of cells from F in wounds created by Incucyte wound-maker. p-value is represented as 2-way ANOVA, Sidak’s multiple test (**H**) Representative images from G. (**I**) Quantification of spheres formed by Calu-1 TS^HIGH^ and TS^LOW^ cells in low adherent cultures at low-seeding density. p-value is Student’s t-Test. (**J**) Quantification of E-CAD and VIM in A549 cells 5 days after sorting in TS^HIGH^ and TS^LOW^ based on the scheme in A. (**K**) Proliferation-independent real-time migration of A549 cells from I in a monolayer wound. p-value is represented as 2-way ANOVA, Sidak’s multiple test (**L**) Western blot of lysates from indicated time points showing reversal of EMT phenotype in A549 TS^HIGH^ and TS^LOW^ cells. E-CAD is a trypsin sensitive membrane cell-adherence protein and could not be detected in sorted cells in suspension on day 0. Error bars represent standard deviation.

Interestingly, a rapid reversion of EMT phenotype was observed in the sorted cells, concomitant with the normalization of *TYMS* promoter activity (**Supp. Figure 2A**). This was more evident in functionally distinct A549 cells sorted for *TYMS* promoter activity (**Supp. Figure 2B)** and recapitulated the EMT phenotype observed in Calu-1 (**Figures 2J-K)**. Sorted cells showed higher TS levels in TS^HIGH^ a day after sorting, followed by a complete normalization of TS and EMT markers after few passages (**Figure 2L**). Therefore, the phenotypic alterations observed between sorted cells were transient and in match with the differences in TS levels. These data strongly indicate a direct control of TS on EMT phenotype and cell plasticity, which was further confirmed by knockdown and overexpression approaches (**Supp. Figure 2C**).

### TS regulates EMT genes in NSCLC

Further, to identify the mediators of TS-promoted EMT, RNA was sequenced from Calu-1 TS^HIGH^ and TS^LOW^ cells in parallel with A549 cells with TS knockdown. Pathway analyses consistently indicated EMT among the topmost differentially regulated pathways (**Figure 3A-B)**, confirming the EMT switch observed with E-CAD and VIM. KRT19, SPARC, SPOCK, LINC00707 (lung cancer promoting lincRNA), FOSL1 and AXL (identified as downstream targets of TS as they appeared in both signatures) were qPCR validated in both cell lines (**Figure 3C, Supp. Figure 3A**). Of these genes, SPARC, FOSL1 and AXL, that have an established role in EMT in NSCLC(17–19), were strongly down regulated at protein level in A549 cells with TS knockdown (**Figure 3D**). Differentially expressed genes (DEGs) were used to derive a knockdown score, which predicted a worse survival associated with lower TS knockdown (higher TS levels, **Figure 3E**) and correlated with published EMT gene signature (**Figure 3F**). This indicated that TS-mediated EMT is empowered with its own prognostic impact, i.e. contributes to the adverse prognosis of NSCLC with high TS levels (Figure 1C-D), suggesting a role for TS beyond proliferation. Several EMT transcription factors were identified from DEGs in endogenously TS^HIGH^ and TS^LOW^ cells, from which siRNA mediated knock-down of HOXC6 and HMGA2 reduced expression of TS in A549 (**Figure 3G**) and Calu-1 (**Supp. Figure 3B**). Thus, these results identified a network of TS mediated EMT, where HOXC6 and HMGA2 are upstream of TS and AXL, SPARC and FOSL1 are downstream mediators (**Figure 3H**).

**Figure 3.**
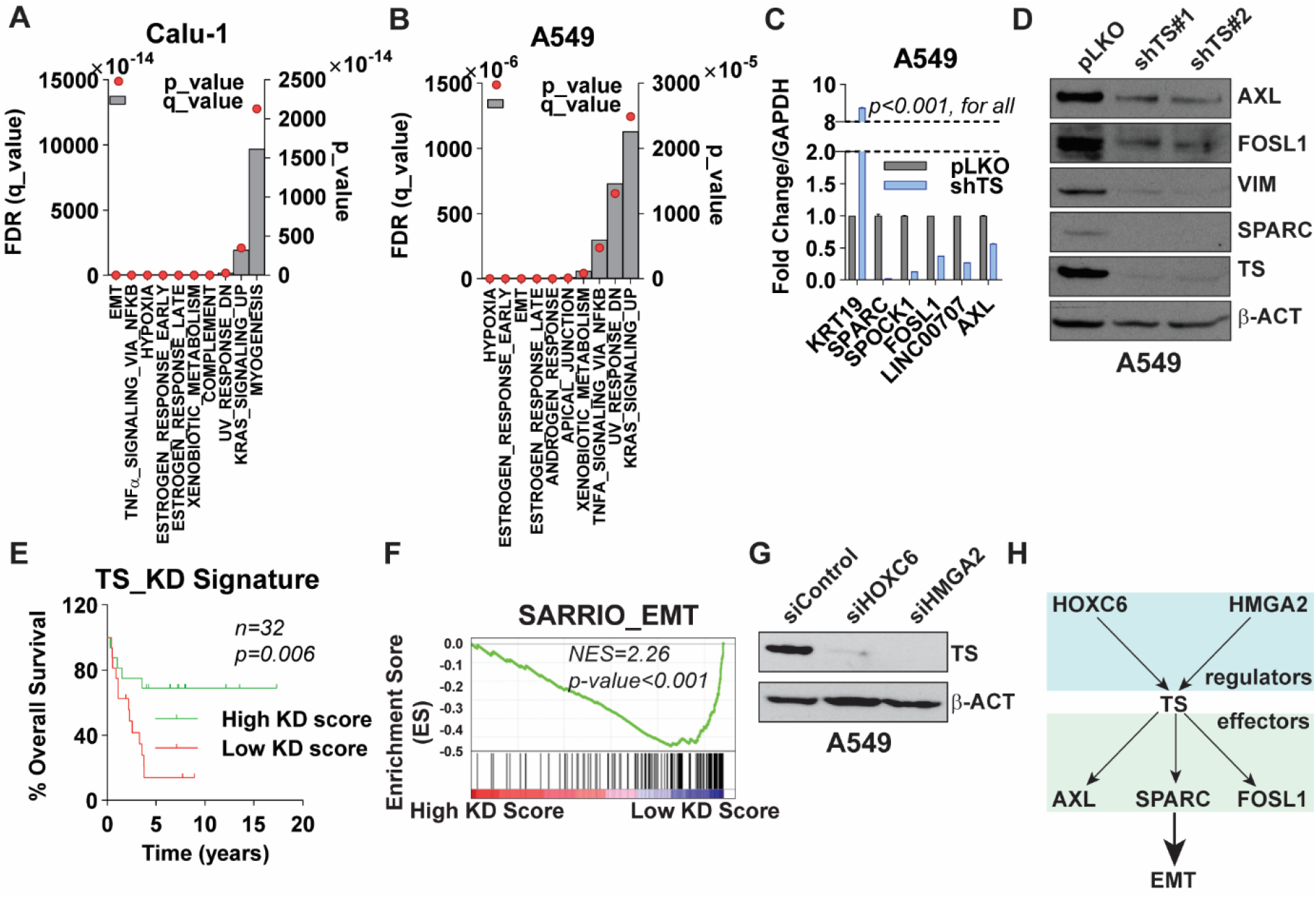
TS expression correlates with EMT gene signature. Gene set enrichment analysis of RNA-seq data of (**A**) Calu-1 cell sorted for high and low TYMS promoter activity and (**B**) A549 cells after transduction of shTS and non-targeting pLKO backbone as control. (**C**) q-RT-PCR quantification of gene signature from A549 cells. p-value represents multiple t-Test (**D**) Validation of TS-mediated downregulation of indicated genes at protein levels in A549 cells with TS knock-down. (**E**) Survival analysis of NSCLC patients from data set GSE101929 based on a knock-down score generated from A549 gene signature in B. p-value is represented as Log-rank (Mantel-Cox) test. (**F**) Gene set enrichment analysis with overlapping of TS knock-down score with EMT-signature in SSC patients from data set GSE4573. (**G**) A549 cells transiently transfected with 50nM siRNA targeting transcription factors HOXC6 and HMGA2 identified from A. (**H**) Scheme showing the regulators (HOXC6, HMGA2) and downstream effectors (AXL, SPARC, FOSL1) involved in TS mediated EMT. Error bars represent standard deviation.

### Depletion of TS mitigates metastasis *in vivo*

Finally, *in vivo* approaches were used to confirm the role of TS on EMT and metastasis. Ts (mouse TS) expression was quantified in morphologically and functionally distinct mesenchymal-like (Ladi 3.1) and epithelial-like (Ladi 2.1) cells (**Supp. Figures 4A-B)**, isolated from the same mouse model of NSCLC (p53^fl/fl-LSL^ KRAS^G12D/+^). Ts positively correlated with Vim and negatively with E-Cad (**Supp. Figure 4C**). Furthermore, to functionally evaluate *in vivo* effects of TS alteration on metastatic colonization, *Tyms* gene was knocked down in murine Lewis lung carcinoma cell line LL/2 using stably transduced shRNA. A moderate Ts depletion (**Figure 4A**) did not affect proliferation (**Supp. Figure 4D**), as we had previously seen that TS needs to reduce beyond a threshold to diminish proliferation(12). Furthermore, Ts knockdown did not hamper growth of primary tumors from the cells subcutaneously injected in flanks of syngeneic mice (**Figure 4B**), confirming our previous observation in triple negative breast cancer and NSCLC cell lines that TS must be reduced below a threshold level to effect proliferation(11, 12). However, when injected in the tail vein, knocked down cells showed a highly significant reduction in lung metastatic colonization (**Figures 4C-D**), and the mice carrying cells with Ts depletion showed a significantly prolonged survival (**Figure 4E**).

**Figure 4.**
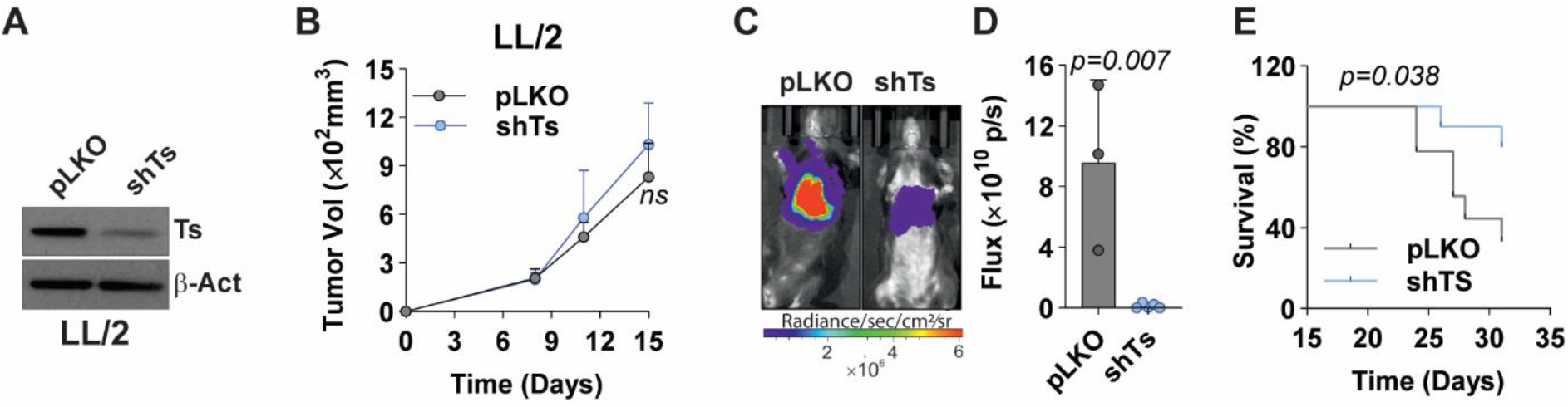
TS expression determines metastatic colonization in vivo. (**A**) Western blot quantification of Ts in murine LL/2 cells. pLKO has been used as a non-targeting control. (**B**) Quantification of primary tumour growth after subcutaneous injection of 1×10^6^ LL/2 cells from H in C57BL/6 mice (n=8). p-value is represented as Student’s t-Test. (**C-D**) Quantification of metastatic localization of luciferase expressing Ts knock-down and pLKO LL/2 cells in lungs of C57BL/6 mice. 5×10^5^ were injected in the tail-vein of mice (n=10) and the luciferase activity was quantified by IVIS as readout for lung metastasis after 4 weeks. (**E**) Survival curves from mice in K. p-value is represented as Log-rank (Mantel-Cox) test.

## Discussion

TS has been widely used as a chemotherapeutic target(20) ascribed to its role in proliferation. This study experimentally validated this concept, as TS was one of the highest ranked target gene identified based on the CRISPR/Cas9 screen analyzed in this study. However, chemotherapeutic drugs that target TS, might also mitigate other detrimental features associated with cancer. We confirmed the multifaceted role of TS proliferation and chemoresistance and found a strong correlation with EMT gene signatures and prognosis, highlighting clinical, as well as biological relevance of TS in NSCLC. Interestingly, presented results clearly corroborate that different functions of TS can operate independently, as observed that TS depletion can mitigate EMT phenotypes without triggering proliferation loss (in LL/2 cells and in other previous settings(12)). This provides a strong rationale to revisit clinical and therapeutic aspects of TS in tumor biology and explore its therapeutic potential beyond proliferation.

Pemetrexed is used as a first line treatment for NSCLC patients(21, 22) albeit response to the drug is remarkably varied. TS expression is an important determinant of sensitivity to pemetrexed(15, 23) and marks the worse clinical outcomes of pemetrexed treatment in NSCLC pateints(24). In agreement to this, SCC generally respond poorly to pemetrexed as compared to ADC(25), partly attributed to higher TS expression in SCC compared to other histological subtypes(14, 26). However, there is also a considerable variability to pemetrexed response with a given histological subtype(27, 28). EMT could be an additional source of variability, as it is a key driver of chemoresistance against pemetrexed in NSCLC(29). In this study we have linked higher TS expression not only to pemetrexed outcome but also to EMT, indicating that TS has an important role in establishing a connection between EMT and pemetrexed resistance. This connection could also be extrapolated to genes that constitute the TS regulatory network, such as FOSL1, which regulates pemetrexed resistance in coordination with EMT-TF ZEB1(18). Therefore, a more inclusive biomarker signature needs to incorporate EMT genes and downstream genes like FOSL1, in addition to TS, for robust prediction of response to pemetrexed in NSCLC(30).

In NSCLC, EMT enhances the inflammatory tumor microenvironment leading to activation of multiple immune checkpoint proteins, including PD-L1.(31) A recent clinical trial has demonstrated a better outcome in NSCLC when pemetrexed is administered in combination with a PD-L1 inhibitor, pembrolizumab(32). Since, TS drives EMT and EMT has been shown to modulate response to immune therapy, a functional correlation between TS expression and susceptibility to immunotherapy could be deduced in NSCLC. In fact, we identified interleukins such IL-6, IL-7 and IL-32 in our TS signatures, which have been previously linked with poor prognosis and metastasis in NSCLC(33–35). Hence, a follow up study is needed to validate this correlation, as the two drugs are frequently combined.

This study also underscores the plasticity of cancer cells with mixed EMT population, as was reported in cells sorted for high and low TS expression (**Figure 2L**). TS expression has been previously shown to be stimulated by chemotherapy, as a cellular defense mechanism(36), and these data add the notion that chemotherapy-induced TS could lead to the adjustment of EMT phenotypes in patients, that, in turns, might influence the efficacy of the treatment. This aspect could be taken in consideration for the implementation of therapeutic strategies combining EMT-suppressing drugs and chemotherapy, or for the future design of the next generation of TS-inhibitors, which should not enhance TS levels.

We furthermore establish the role of TS in metastasis, where TS knock-down abrogated the metastatic colonization and improves mice survival without affecting proliferation and growth of primary tumor. This observation indicate that TS strongly influences the success against selection pressure at the metastatic site. Further retrospective validation in patients can establish TS as a metastasis marker in NSCLC.

Finally, this study provides a perspective for a network that could integrate different signaling pathways to effectuate various aspects of cancer progression that are mediated by TS (**Supp. Figure 3G**), worth further investigation. Different transcriptional regulators and effector proteins identified in this study have an established role in EMT in NSCLC and connect with master EMT-TFs. HMGA2, for instance, affects proliferation and metastasis by regulating TWIST(37), FOSL1 regulates chemotherapy, exogenous SPARC promotes invasion and metastasis by activating SNAI1(17, 37) and AXL activates TWIST to affect cell cycle(19). A follow up study could further substantiate TS as an integration point for these pathways resulting in a cumulative readout in terms of metastasis.

Thus, this study provides strong evidence that TS, apart from proliferation marker, also regulates EMT in NSCLC. Targeting EMT-related processes could represent a promising therapeutic strategy to suppress the aggressiveness of TS-overexpressing NSCLC.

## Supporting information

Differentially expressed genes

## List of Abbreviations

TS: thymidylate synthase (human)
Ts: thymidylate synthase (mouse)
EMT: Epithelial-mesenchymal transition
NSCLC: non-small cell lung cancer.

## Ethics Approval

The use of retrospective solid tumor tissues for the immunohistochemical and qPCR study was approved by the Research Ethics Committee of the San Luigi Hospital/University of Turin (approvals n.167/2015 and 204/2016). In vivo experiments were performed by skilled experimenters trained according to FELASA guidelines. Animal protocols were approved by the Institutional Animal Care and Use Committee of the Regierung von Unterfranken.

## Authors’ contributions

**MAS** was involved in conception, design, acquisition, and analysis of *in vitro* experimental data and drafting of the manuscript. **PNG** was involved in acquisition and analysis of *in vivo* experimental data. **VR** was involved in data acquisition and bioinformatic analysis. **BP, AS and MEV** were involved in *in* vitro data acquisition, **RN** performed RNA-sequencing, **OS** performed analysis of RNA-Seq data, **IR** and **PB** acquired patient data, **HS** helped in design of the *in vivo* work, **IAA** helped with interpretation of RNA-seq Data, **OSa** helped with generation and interpretation of knock down score and critical revision of the manuscript, **MV** was involved in interpretation of patient data and critical revision of the manuscript, **PC** was involved in project supervision, experimental conceptualization and design, analysis and interpretation of data and drafting of manuscript.

All authors read and approved the final manuscript.

## Acknowledgments

Authors would like to thank Dr. Markus Diefenbacher, Dept. of Biochemistry and Molecular Biology, University of Würzburg for providing Ladi 2.1 and Ladi 3.1 cells and Prof. Dr. Susetta Finotto, Dept. of Molecular Pneumology, University Hospital Erlangen, for LL/2 cells. The sorting was performed at Core Unit for Cell Sorting and Immunomonitoring at Nikolaus Fiebiger Zentrum, Friedrich-Alexander University of Erlangen Nuremberg. Results were partially presented at the 18th World Conference on Lung Cancer in Yokohama, Japan.

## Supplementary figures

**Supplementary Figure 1. TS is an essential NSCLC gene with prognostic/predictive power and correlates with EMT signature**. (**A**) Graph showing *TYMS* priority score in a pan-cancer analysis. (**B-C**) Predicted overall survival in non-small cell lung cancer (NSCLC) patients separated according to the expression level of TS in mentioned datasets. P-value is Logrank test. (**D**) NSCLC cell lines arranged in epithelial-like, intermediate and mesenchymal-like phenotype bases on the ratio of *VIM/CDH1*. (**E-F**) Gene-set enrichment analysis of indicated datasets with high and low TS correlating with EMT gene signature in GEO dataset GSE72094 and in TCGA lung adenocarcinoma expression dataset, respectively.

**Supplementary Figure 2. Endogenous expression of TS is determinant of EMT phenotype**. (**A**) FACS quantification of Calu-1 cells showing normalization of TS promoter activity. (**B**) FACS plots depicting the separation of high and low population of A549 cells sorter for *TYMS* promoter activity. The numbers on the FACS plot represent the percentage of TS^HIGH^ and TS^LOW^ population in the unsorted cells. (**C**) Quantification of TS and E-CAD in Calu-1 cells with shRNA mediated knock-down and overexpression of TS. pLKO and empty vector have been used as controls.

**Supplementary Figure 3. Upstream and downstream genes were identified in NSCLC**. (**A**) mRNA quantification of genes identified from RNA-seq overlapping in Calu-1 TS^HIGH^ and TS^LOW^ cells and A549 shTS cells in Calu-1. p-value represents multiple t-Test. (**B**) siRNA mediated knock-down of specified transcription factors in Calu-1 cells.

**Supplementary Figure 4. TS expression determines Metastasis in vivo**. (**A**) Pictures showing morphological distinction in Ladi cells isolated from a NSCLC mouse model. Ladi 2.1 is a cell line with epithelial trait whereas Ladi 3.1 cells harbour mesenchymal features. (**B**) Real-time migration quantifying migration difference between Ladi 2.1 and Ladi 3.1 cells (**C**) Western blot showing difference in Ts and EMT markers in Ladi cells. (**D**) Graph comparing proliferation rates between LL/2 shTs and pLKO cells. p-value is represented as 2-way ANOVA, Sidak’s multiple test. Error bars represent standard deviation. (**E**) A perspective hypothesis depicting potential upstream regulator and effectors of TS mediated EMT and their possible integration. HMGA2 and HOXC6 upregulate the expression of TS, where TS modulates the expression of proteins (AXL, FOSL1 and SPARC) that have been shown to have a role in EMT. Pathways and the responses elicited by each protein have been depicted in a different colour.

