## Supplementary figures and images for "Thymidylate synthase drives the phenotypes of epithelial-to-mesenchymal transition in non-small cell lung cancer"

### Supp. Figures

**Supp. Figure_1**

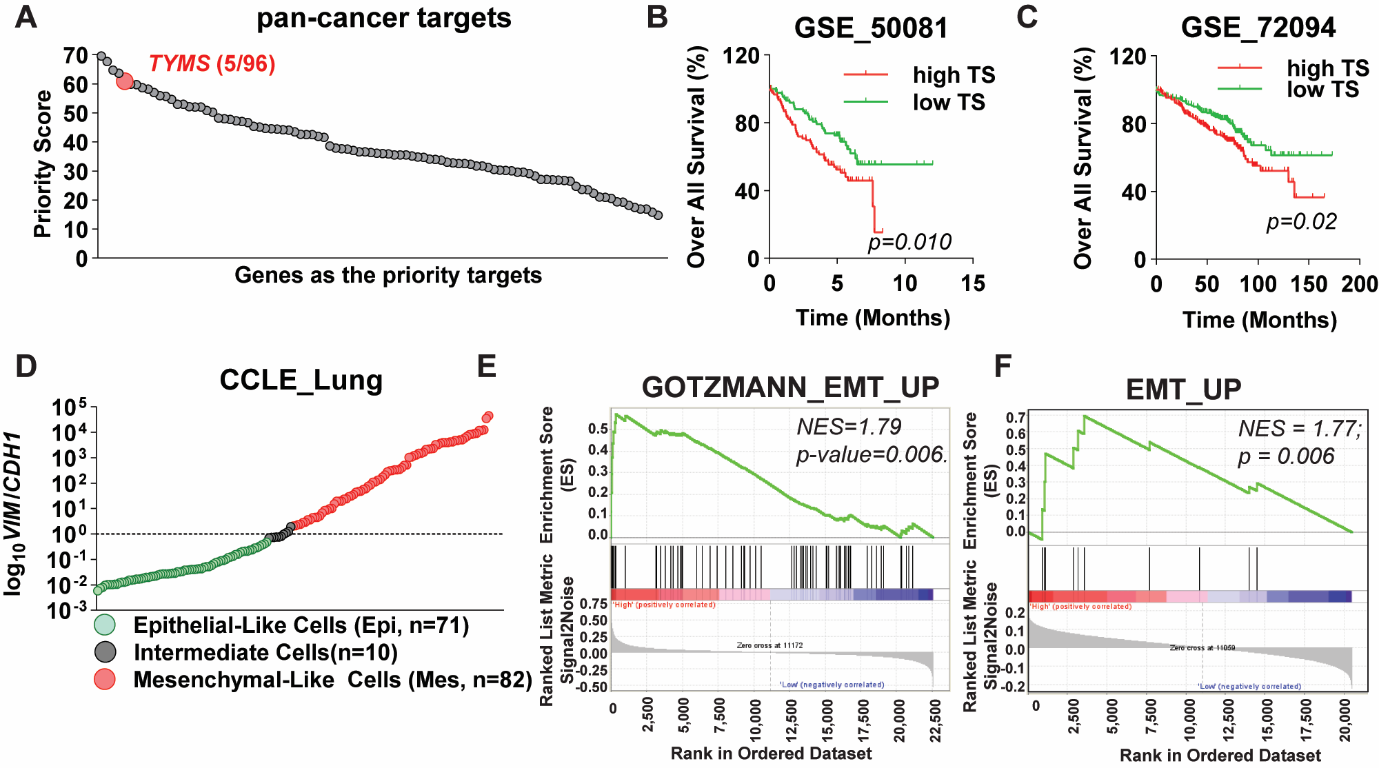

**Supp. Figure_2**

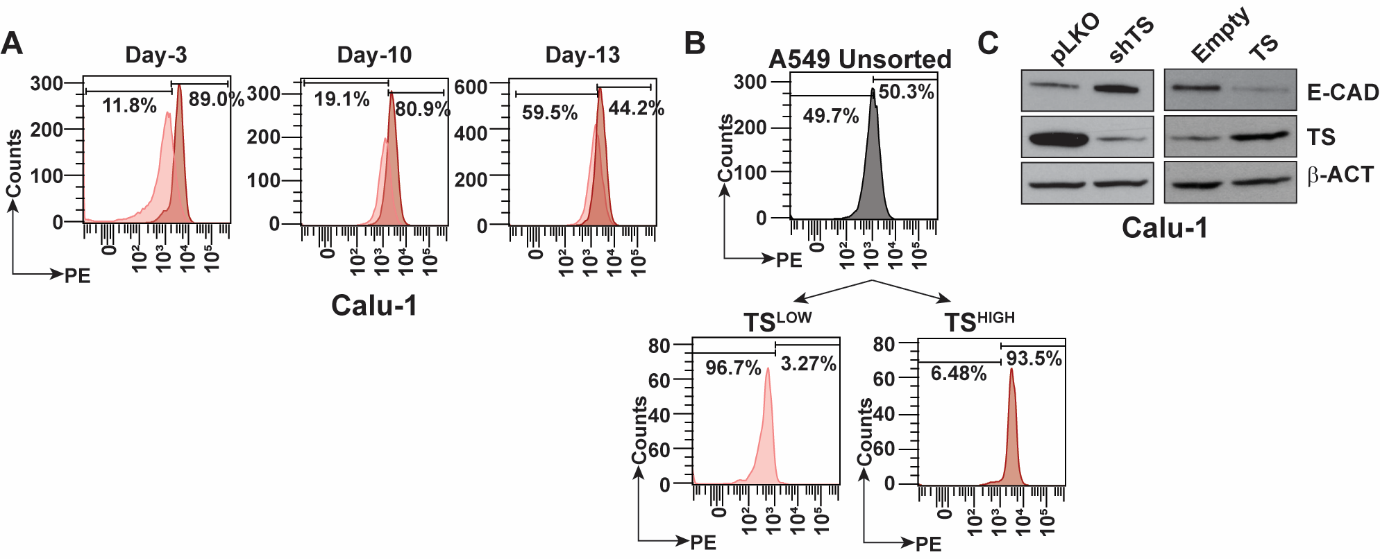

**Supp. Figure_3**

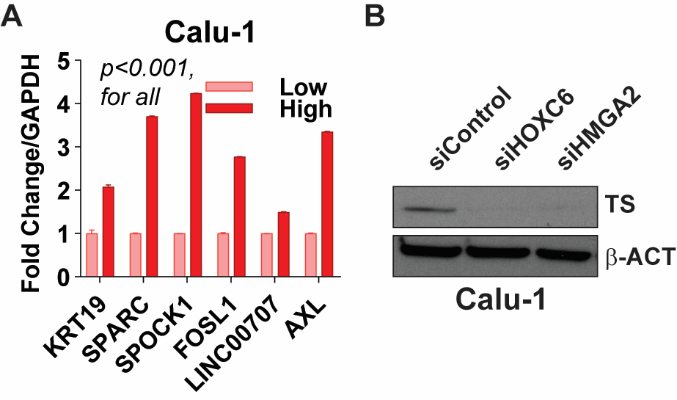

**Supp. Figure_4**

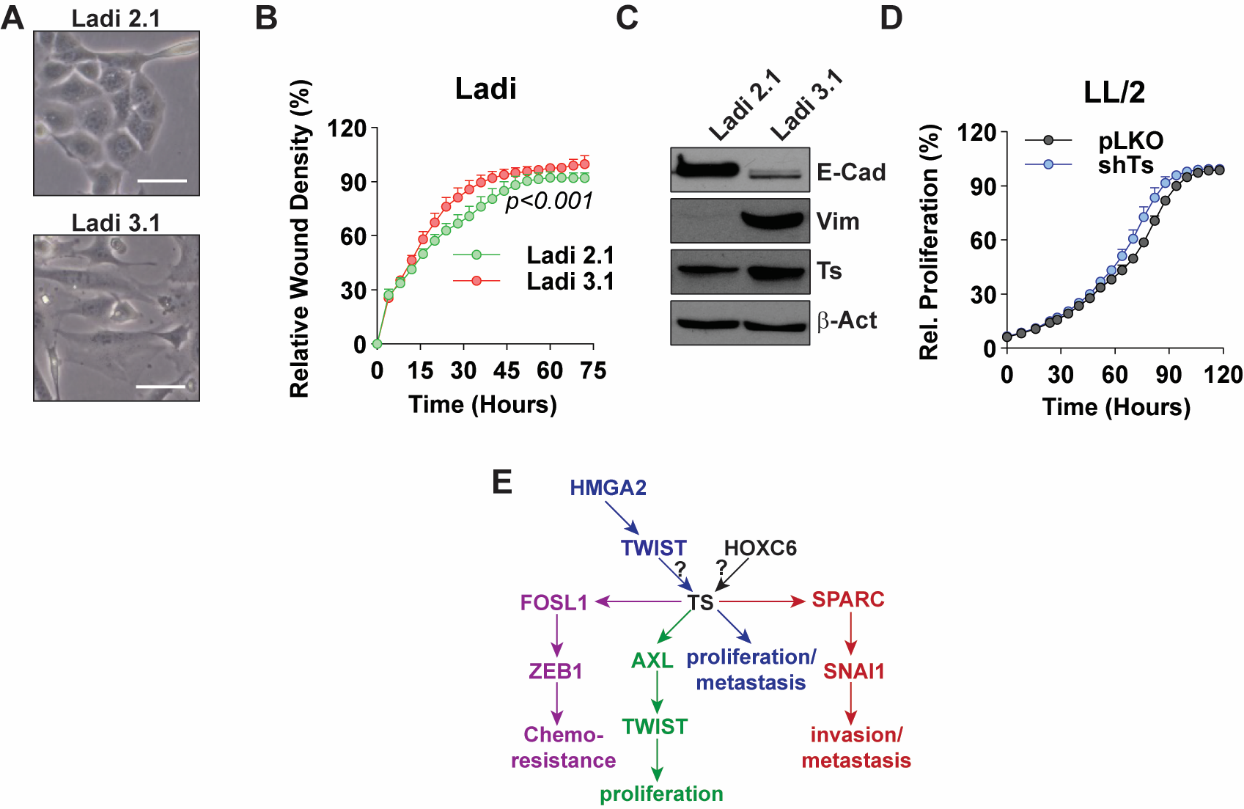
